## Supporting Information - Figures for "Effector Innovation in Genome-Reduced Phytoplasmas and Other Host-Dependent Mollicutes"

**This PDF file includes:**

Supplemental Figures S1 to S17

**Other Supplementary Materials for this manuscript include the following:**

Tables S1 to S11 (Tables_PhAMES - Excell sheet)

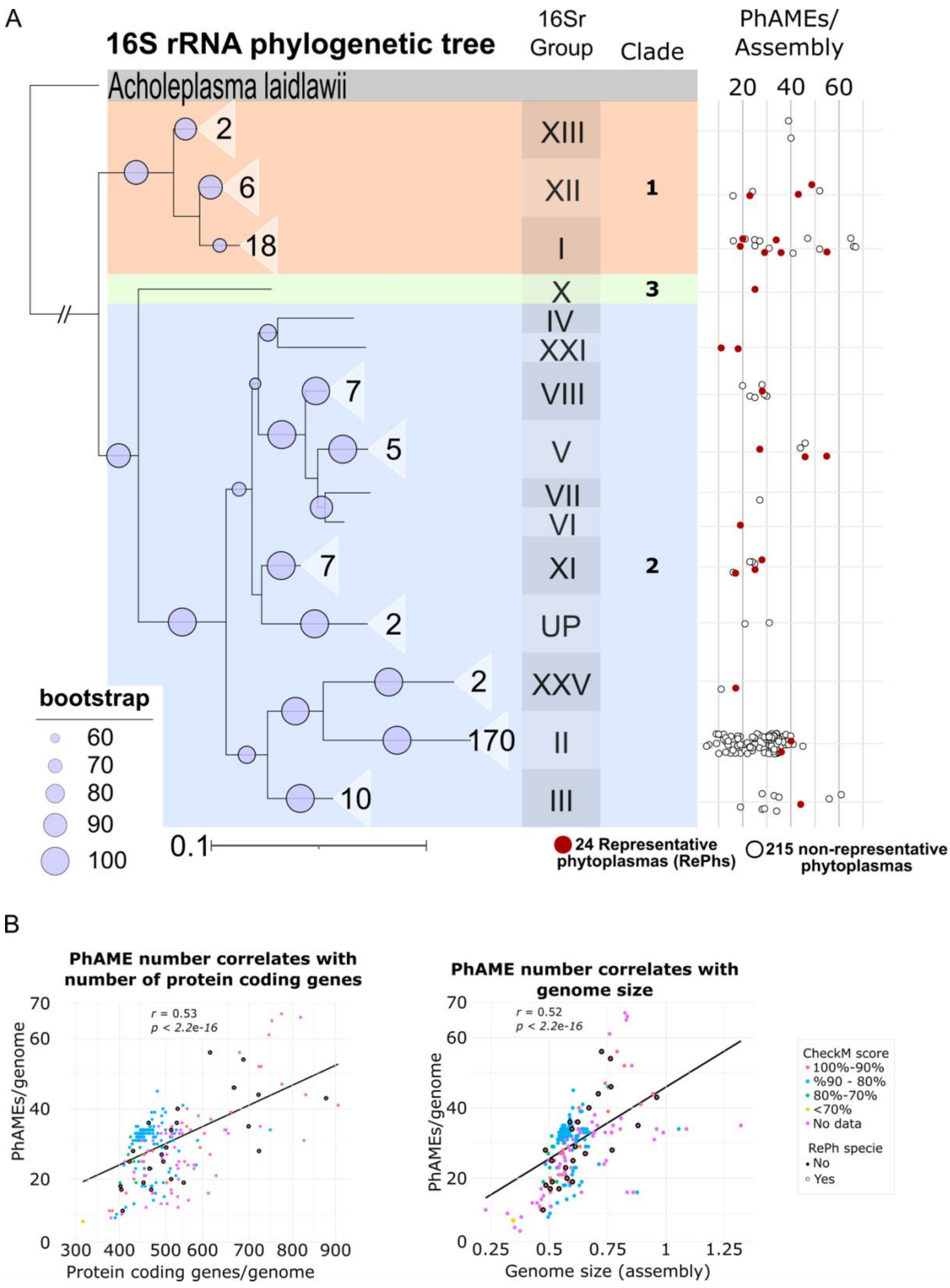


**S1 Figure. Phytoplasmas contain a variable number of PhAMEs.** (**A**) A maximum likelihood phylogenetic tree was constructed using 16S rRNA gene sequences of ≥ 98% completeness aligned with MUSCLE (101) and inferred with RAxML using the GTRGAMMA model and 100 bootstrap replicates. Tree representation was done with iTOL. Three accessions belonging to the 16Sr IX group (Pigeon pea witches'-broom group) and two unclassified phytoplasmas were not included in the tree. *Acholeplasma laidawii* 16S rRNA gene was used as outgroup. Bootstrap values ≥ 60% are indicated by purple circles. Branch lengths are proportional to the nucleotide substitution rate, see scale bar for reference. The phylogenetic tree highlights three phytoplasma groups, indicated by distinct colours, and the separate 16Sr groups indicated with Roman numbers. 'UP' indicates unclassified phytoplasmas of the 16Sr group. For branches containing more than one species, the number of species within each branch is specified in triangles. The triangle numbers include phytoplasmas with < 98% 16S rRNA gene sequence completeness, but for which the 16Sr group is known. The PhAMEs/genome for each species is shown to the right of the phylogenetic tree, with PhAME numbers from the initial 24 phytoplasmas shown in red and those belonging to the 215 in white (See Fig. 1A). (**B**) Positive correlation between the number of PhAMEs and both the total number of protein-coding genes (left) and genome size (right). Point colour represents the CheckM score, an estimate of genome completeness (57). Genomes from representative phytoplasmas (RePh) are outlined in black. Pearson correlation coefficient (*r*) and p-value (p) are shown.

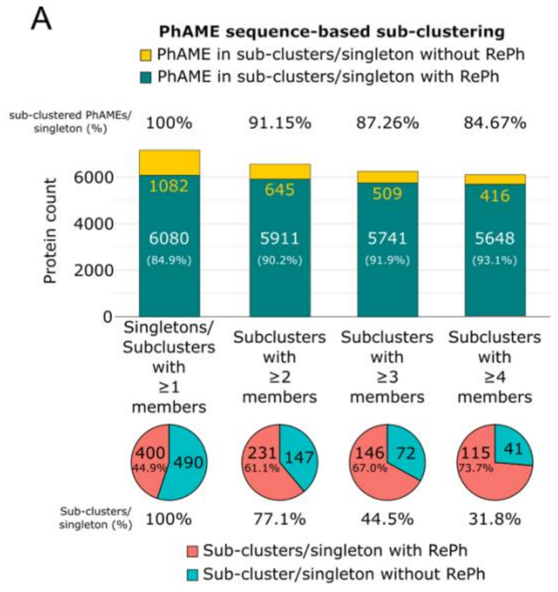


**S2 Figure. Most PhAMEs share sequence similarity with effectors from representative phytoplasma (RePh) species.** Top: Number of PhAMEs grouped by sequence-based sub-cluster size. Green bars indicate PhAMEs that cluster with those from RePh species; yellow bars indicate PhAMEs that do not. Percentages above bars reflect each group's proportion of the total PhAME set. PhAMEs with mature lengths < 13 amino acids are included in the "PhAMEs in sub-clusters/singletons without RePh" category. Bottom: Number of sub-clusters and singletons grouped by membership size. Salmon area denotes sub-clusters containing at least one RePh-derived PhAME; cyan area denotes those without. Percentages below pie charts represent the proportion of sub-clusters in each category relative to the total.

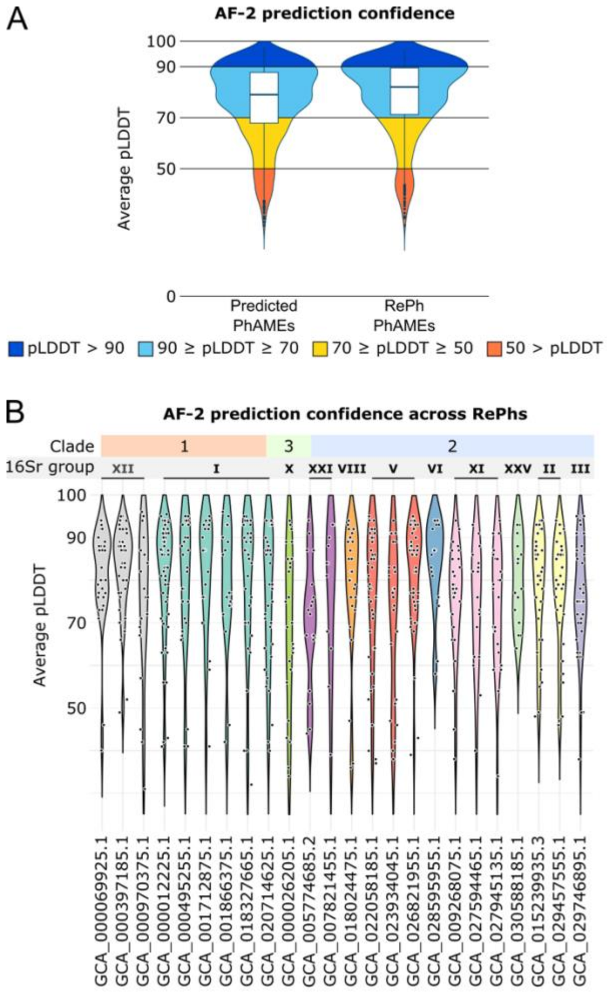


**S3 Figure. Most PhAME structures are predicted at high confidence.** (**A**) Distribution of average pLDDT scores for AlphaFold v2.0-(AF2)predicted models of all PhAMEs and those from the RePh species subset only. Box plots represent the interquartile range (IQR), with whiskers indicating the full range (minimum to maximum). Individual outliers are shown as jittered points. (**B**) Average pLDDT scores of PhAMEs in each of the 24 representative phytoplasma (RePh) species. The three major phytoplasma clades and their corresponding 16Sr groups are indicated at the top of the graph.

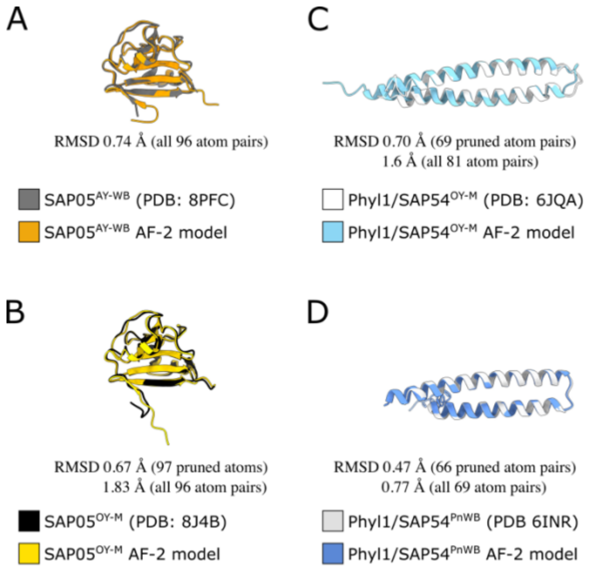


**S4 Figure. AF2-predicted models match the experimentally determined crystal structures of the SAP05 and Phyl1/SAP54 PhAMEs.** Structural alignment of experimentally determined structures of SAP05 from (**A**) AY-WB and (**B**) OY-M phytoplasmas and SAP54 from (**C**) OY-M and (**D**) PnWB phytoplasmas with their corresponding AlphaFold-2 models. RMSD scores below the aligned structures were calculated by the Matchmaker command from ChimeraX.

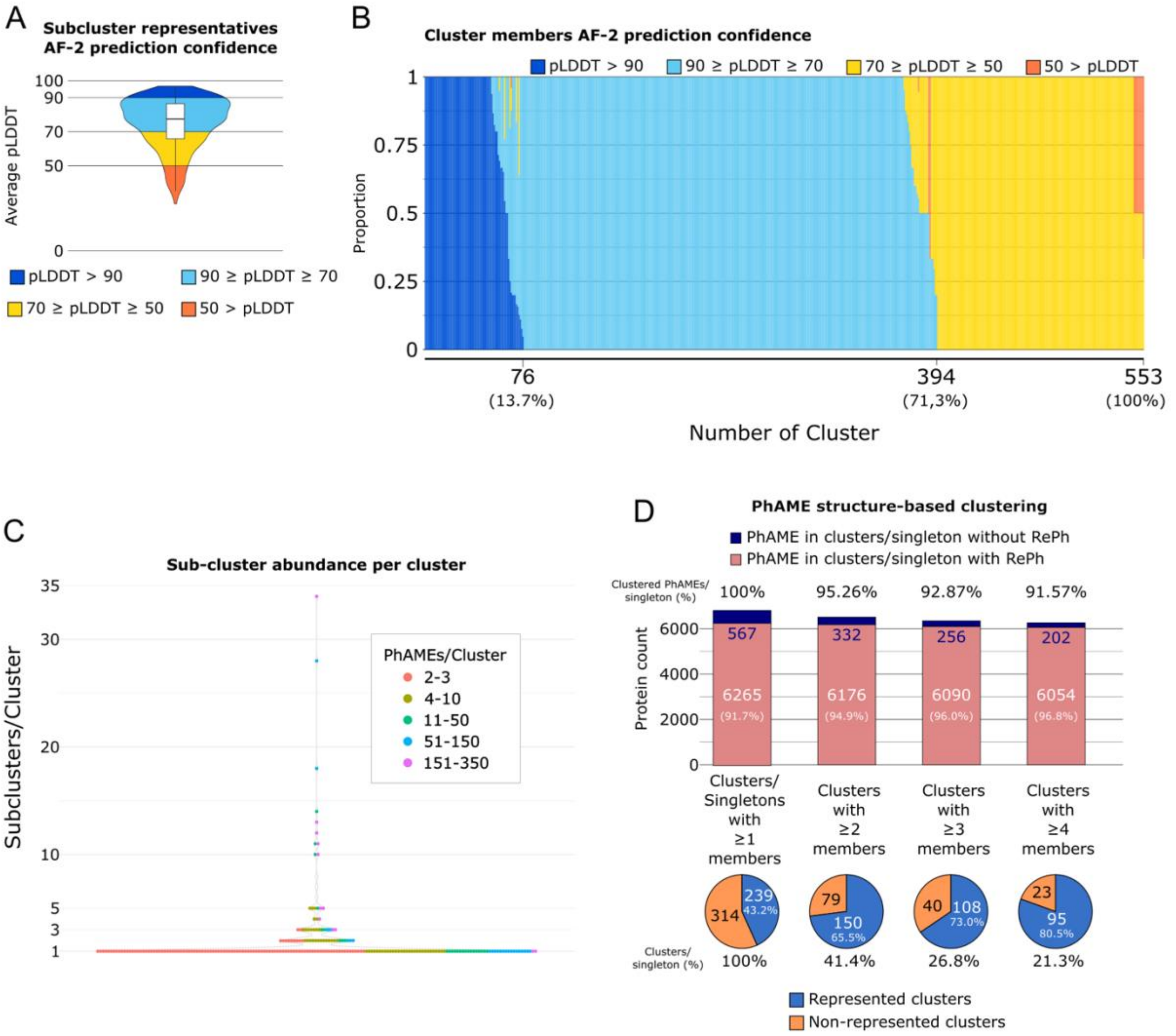


**S5 Figure. Most PhAMEs are grouped into structurally related clusters, supported by high-confidence structural models.** (**A**) Average pLDDT score distribution of AlphaFold-2-predicted models for sub-cluster representatives, where each representative is the sequence with the highest average pLDDT score in its sub-cluster. Box plots represent the interquartile range (IQR), with whiskers indicating the full range (minimum to maximum). (**B**) Proportion of confidence prediction categories across clusters and singletons ordered by the highest proportion of members falling into their most confident prediction category. X-axis labels indicate the number of clusters containing at least one member within the corresponding confidence category. (**C**) Number of sequence-similarity-based sub-clusters within structure-based clusters. Number of PhAMEs with membership to each cluster is indicated in colours. Singletons were not included. (**D**) Top: Number of PhAMEs assigned to structure-based clusters, grouped by cluster size. Salmon bars represent PhAMEs that cluster with those from RePh species, while dark blue bars represent PhAMEs that do not. Percentages above bars are calculated relatively to the total number of clustered PhAMEs. Percentages within bars are related to PhAMEs with assigned clusters. Bottom: Number of structure-based clusters or singletons, grouped by membership size. Blue area denotes clusters containing at least one RePh-derived PhAME; orange area denotes those without. Percentages below pie charts represent the proportion of clusters in each category relative to the total.

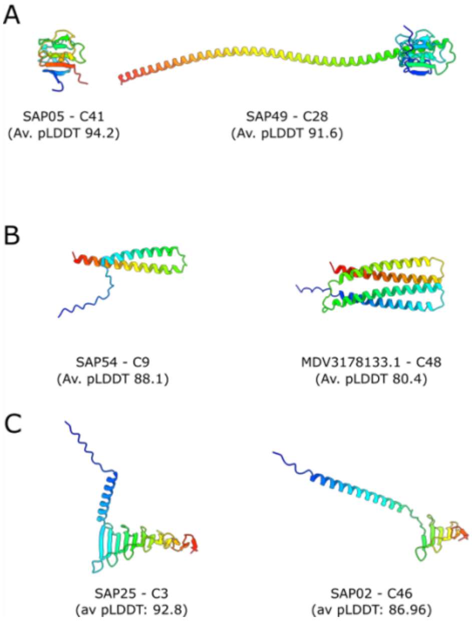


**S6 Figure. Predicted structures of PhAMEs belonging to different clusters with structural similarities. (A)** Structural prediction of SAP05 (left), with membership to cluster C41 and the structurally related SAP49 with membership to C41 (right), both from AY-WB phytoplasma. **(B)** Structural prediction of the AY-WB phytoplasma effector SAP54 (left), a member of the large cluster C9, which comprises numerous structurally similar effectors and is linked to other related clusters such as C48. Shown on the right is the predicted structure of MD3178133.1 from Sweet Potato Little Leaf Phytoplasma, a representative of cluster C48. **(C)** Predicted structures of AY-WB phytoplasma effectors SAP25 (cluster C3, left) and the structurally related SAP02 (cluster C46, right).

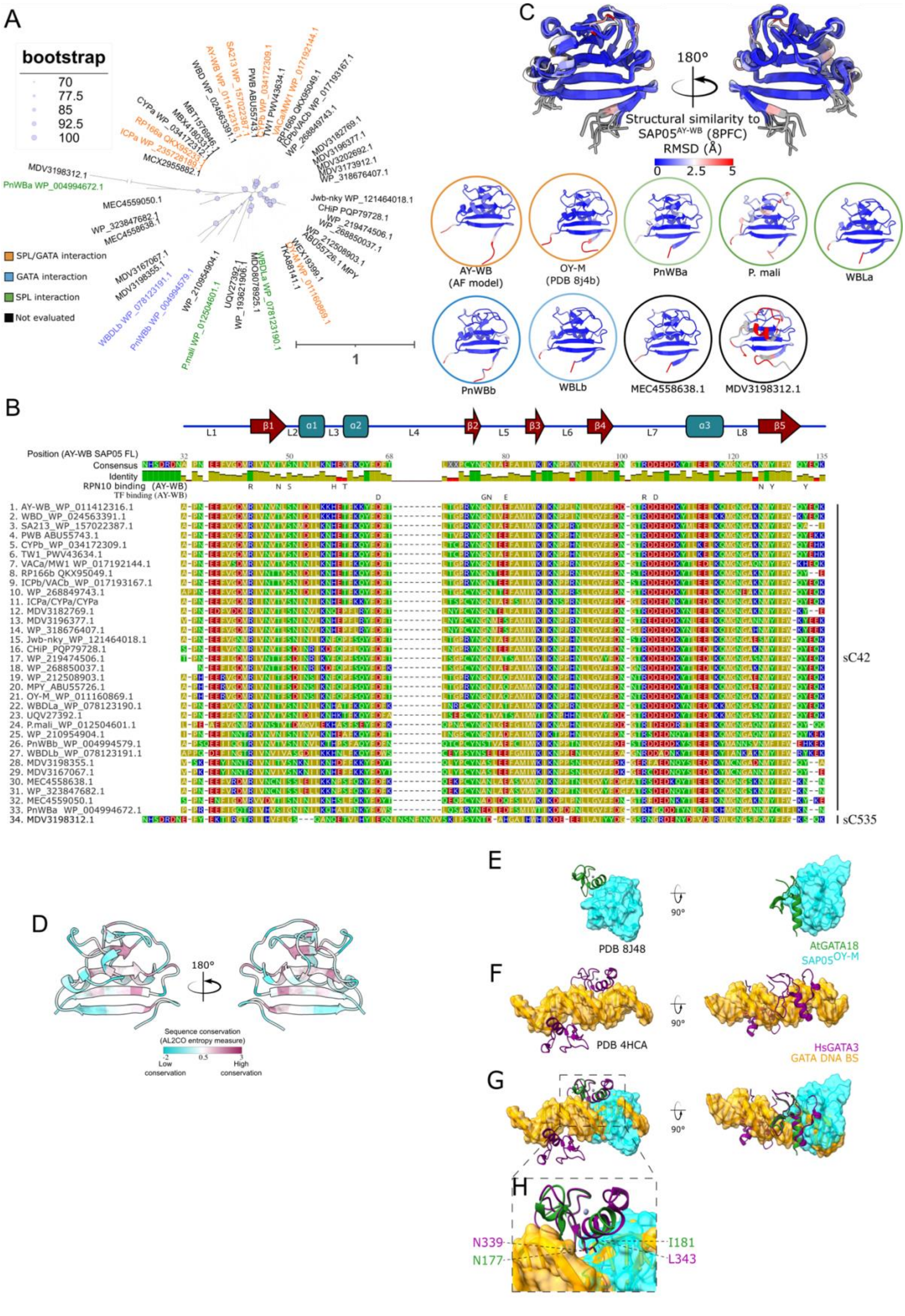


**S7 Figure. SAP05 homologues are structurally conserved.** (**A**) RAxML phylogenetic tree constructed using the maximum likelihood method with mature effector sequences from cluster 41, with 1000 bootstrap replications. Homologues reported in Huang et al., 2021 that failed to be identified by our pipeline were included (30) Bootstrap values ≥ 70% are indicated by purple circles. SAP05 homologs with experimentally validated abilities to mediate degradation of SPL (green), GATA (blue), or both families of TFs (orange) are indicated (30). Branch lengths are proportional to the amino acid substitution rate, see scale bar for reference. Tree representation was done using iTOL web server (135). (**B**) MUSCLE (101) protein sequence alignment of the effectors shown in the phylogenetic tree (panel A). Secondary structure is depicted on top of the alignment using the SAP05$^{AY\text{-}WB}$ full length protein as a reference. Experimentally validated residues of SAP05$^{AY\text{-}WB}$ involved in the interaction with TF or RPN10 are indicated below the consensus sequence of the alignment (black letters). Residues in the alignment are coloured based on their chemical properties. All SAP05 homologs clustered in subgroup sC42, except for one that clustered in sC535, as indicated. Duplicated sequences were included once. **(C**) Top: Superimposition of predicted structures from a select number of SAP05 homologs to the crystal structure of SAP05$^{AY\text{-}WB}$ (PDB: 8PFC), performed using the Matchmaker command in ChimeraX. Structures are coloured by root mean square deviation (RMSD), with blue indicating high similarity and red indicating low similarity to SAP05$^{AY\text{-}WB}$, and regions with RMSD > 5 Å indicated in grey. Bottom: AF2-predicted structural models the same SAP05 homologs with the additional inclusion of MDV3198312.1. Coloured outlines indicate experimentally validated ability to mediate degradation of SPL (green), GATA (blue), or both transcription factor families (orange) (30). **(D**) Level of conservation of residues among SAP05 homologs in the cC42 sub-cluster (excluding MDV3198312.1) mapped onto the SAP05 crystal structure. Sequence conservation was calculated using the entropy-based method AL2CO (118) implemented and visualized in ChimeraX. **(E**) Front (left) and top (right) views of *A. thaliana* GATA18 bound to SAP05$^{OY\text{-}M}$ (PDB: 8J48). **(F**) Front (left) and top (right) views of *Homo sapiens* GATA3 (HsGATA3) binding to its target DNA sequence (GATA DNA BS) (PDB: 4HCA). **(G-H**) Overlapping GATA transcription factors surfaces interact with SAP05 and DNA. Structural superimposition of *A. thaliana* GATA18 in complex with SAP05 (PDB: 8J48) and *H. sapiens* GATA3 bound to DNA (PDB: 4HCA), performed using the Matchmaker command in ChimeraX (G). Close-up view of GATA surfaces that interact with SAP05 and DNA with amino acid residues involved in the interaction with both SAP05 and DNA represented as sticks and labelled (H). Grey spheres indicate $Zn^{2+}$ ions (E-H).

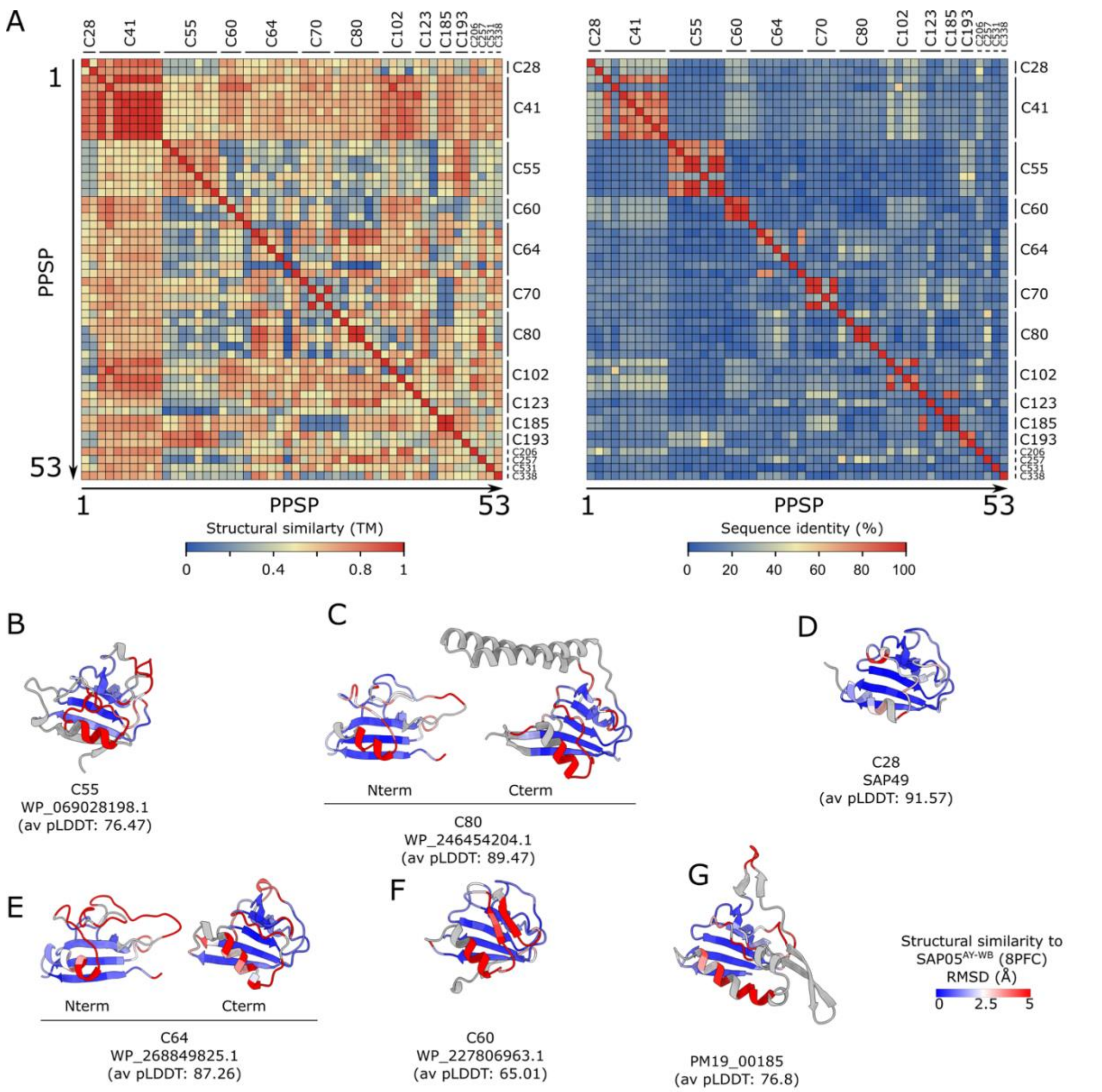


**S8 Figure. SAP05-like proteins share high structural similarity despite low sequence identities. (A)** Levels of structural (left) and sequence (right) similarity among proteins with SAP05-like folds in structure-related clusters. Sequence identity values were calculated based on structure-based alignments using Foldseek. **(B-G)** Parts of proteins with predicted SAP05-like folds from representatives of selected clusters shown in Figs. 2C–F and 2I. Colours indicate structural similarity based on RMSD relatively to the experimentally determined structure of SAP05$^{AY-WB}$, using the Matchmaker command in ChimeraX. Regions in grey indicate RMSD > 5 Å. The representative model with the highest average pLDDT score of each cluster was used for alignment. SAP05-like domains were defined via sequence-independent multiple structure alignment using FoldMason (117), with additional residues included to encompass the full β5 β-sheet. Cluster number and average pLDDT of full-length proteins are indicated below the structures.

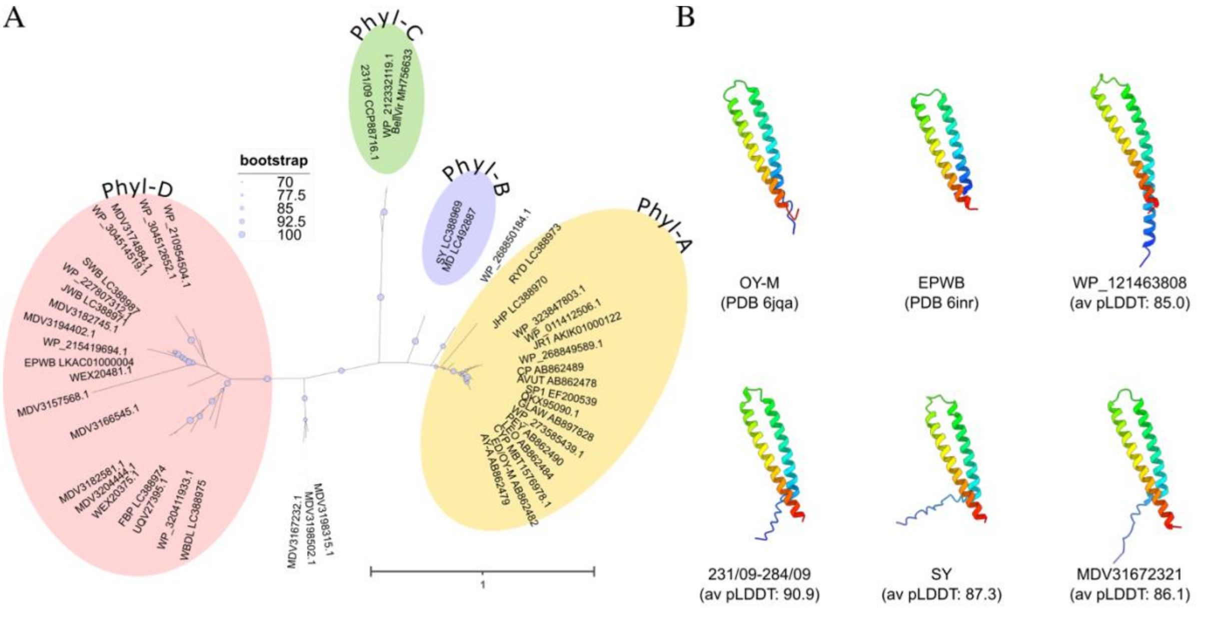


**S9 Figure. SAP54/Phyl1 phylogeny.** (**A**) Maximum likelihood phylogenetic tree constructed from C9 cluster SAP54/Phyl1 effector alignment shown in A with 1,000 bootstrap replicates (IQ-TREE2, (103)). Bootstrap values ≥ 70% are indicated by purple circles. Branch lengths are proportional to the amino acid substitution rate (see scale bar). The tree was visualised using the iTOL web server. Phylogenetic clades identified by Iwabuchi et al. (2020) (70) are colour-coded. (**B**) Crystal structures and AlphaFold-2 models of representative members of the C9 cluster.

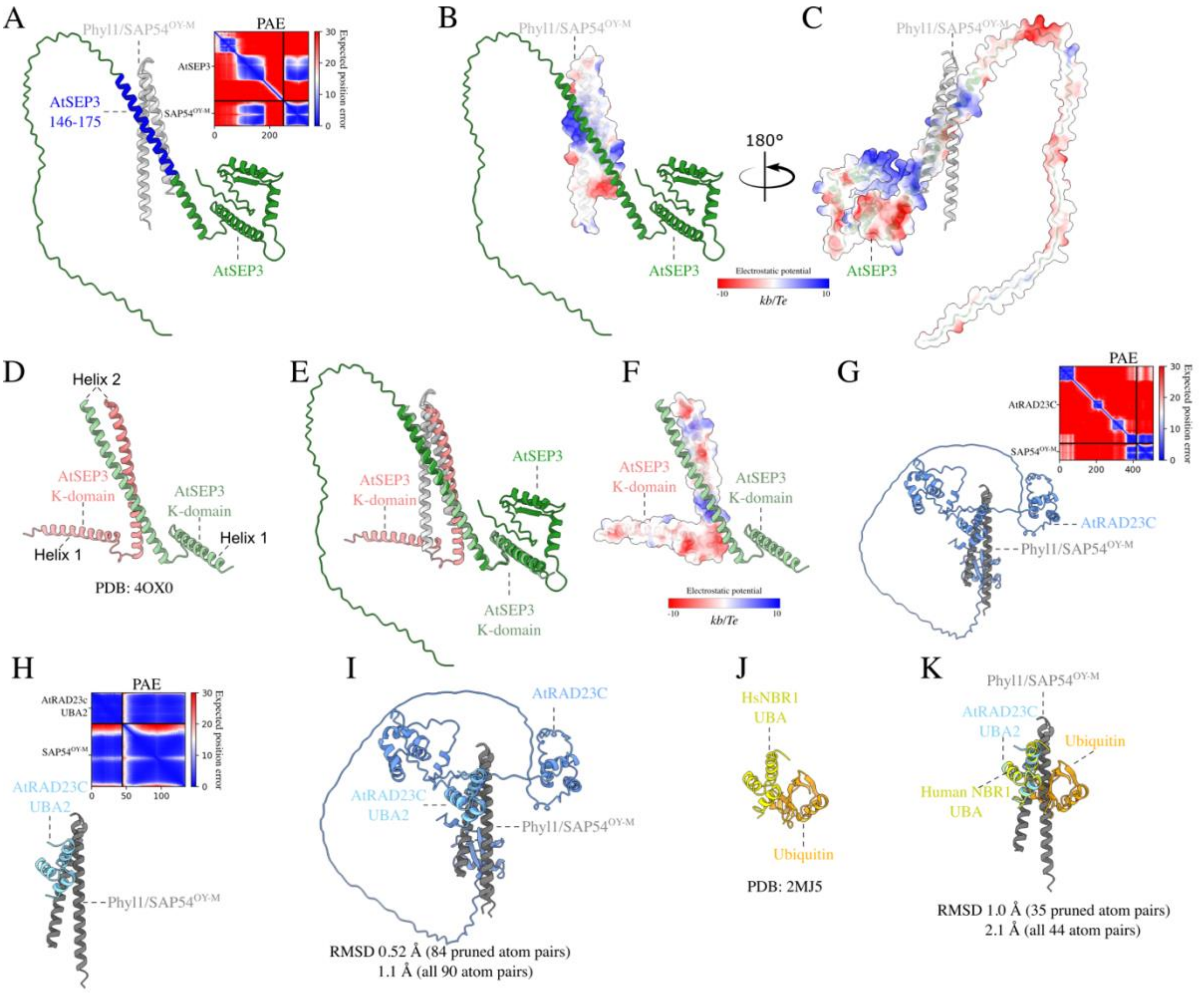


**S10 Figure. Predicted structural basis for the SEP3–SAP54–RAD23 complex.** (**A**) AlphaFold-Multimer (AF-M) prediction of the Phyl1/SAP54$^{OY\text{-}M}$–AtSEP3 complex. AF-M correctly identifies residues 146–175 of AtSEP3 (highlighted in blue) as involved in the interaction with SAP54, consistent with prior data (43). Right: Predicted Aligned Error (PAE) plot for the highest-confidence model, indicating estimated positional uncertainty between residue pairs. Blue represents low predicted error (high confidence), red indicates high error (low confidence). (**B–C**) Surface representations of the predicted SAP54$^{OY\text{-}M}$/Phyl1–AtSEP3 complex, with Coulombic electrostatic potential mapped onto (B) SAP54$^{OY\text{-}M}$/Phyl1 and (C) AtSEP3. Surfaces are coloured from positive (blue) to negative (red) charge. Electrostatic potential was calculated using ChimeraX. (**D**) Crystal structure of the AtSEP3 K-domain dimer (PDB: 4OX0). Helix 1, involved in the dimerization and Helix 2, involved in the dimerization and tetramerization of SEP3 are indicated. (**E**) Structural superposition of the predicted SAP54$^{OY\text{-}M}$/Phyl1–AtSEP3 complex with the AtSEP3 K-domain dimer, highlighting overlapping interfaces involved in MADS-box multimerization and SAP54 binding. (**F**) AtSEP3 K domain with one monomer coloured as in (B). (**G**) Predicted SAP54$^{OY\text{-}M}$/Phyl1–AtRAD23C complex, generated using AF-M. The C-terminal UBA2 domain of RAD23C is correctly predicted to mediate the interaction, as supported by experimental data (Fig. 3A; (43)). (**H**) PAE plot of the highest-confidence SAP54$^{OY\text{-}M}$/Phyl1–UBA2(AtRAD23C) model, coloured as in (B). (**I**) Structural superimposition of the SAP54$^{OY\text{-}M}$/Phyl1–AtRAD23C complex using ChimeraX MatchMaker tool. (**J**) Crystal structure of the UBA domain of human NBR1 in complex with ubiquitin (PDB: 2MJ5). (**K**) Structural superposition of the predicted SAP54$^{OY\text{-}M}$/Phyl1–UBA2(AtRAD23C) complex with the UBA(NBR1)–ubiquitin complex, suggesting that SAP54 targets the ubiquitin-interacting surface of UBA domains.

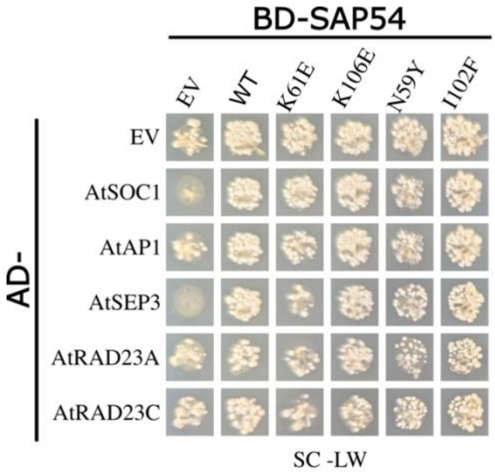


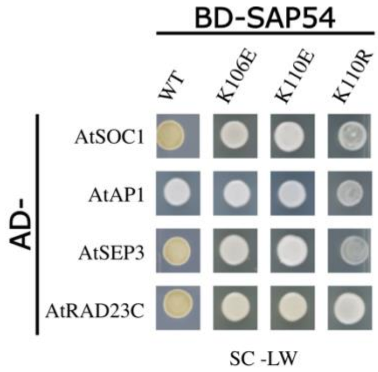


**S11 Figure. Yeast transformation controls for Y2H assays related to Figure 3E**. Growth on double dropout (SC-LW) medium confirms the presence of both constructs. EV: empty vector; AD: GAL4 activation domain; BD: GAL4 DNA-binding domain.

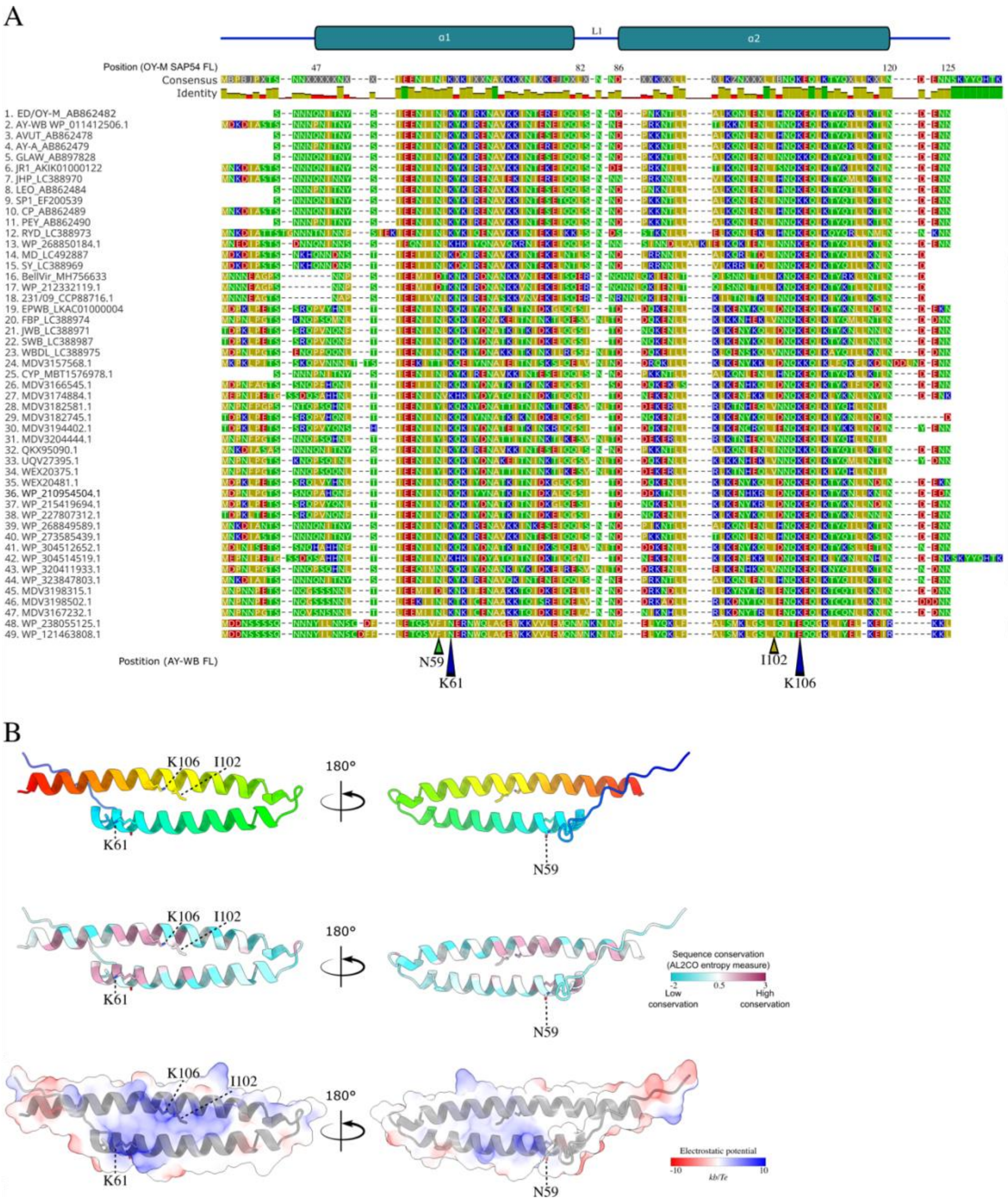


**S12 Figure. SAP54/Phyl1 effectors share conserved residues involved in interactions with plant targets.** (**A**) MUSCLE (101) protein sequence alignment of the C9 cluster SAP54/Phyl1 effectors. Secondary structure is depicted on top of the alignment using the SAP54$^{AY\text{-}WB}$ full length protein as a reference. Their corresponding phylogenetic clades(70) (Phyl-A, Phyl-B, Phyl-C, Phyl-D) are indicated in black on the right. Residues tested for their role in interactions with SAP54 binders (Figure 3) are marked with triangles below the alignment (blue: RAD23 interaction; green: TF interaction). The I102 residue important for maintaining the SAP54/Phyl1 structure is indicated with a yellow triangle. (**B**) Predicted structure of SAP54$^{AY\text{-}WB}$ showing residue conservation and Coulombic electrostatic potential, with the latter mapped onto the protein surface. Sequence conservation was calculated using the entropy-based method in AL2CO (118), and electrostatic potential was computed using the ChimeraX Coulombic command. Residues identified as affecting the interaction between SAP54 and its binders are highlighted.

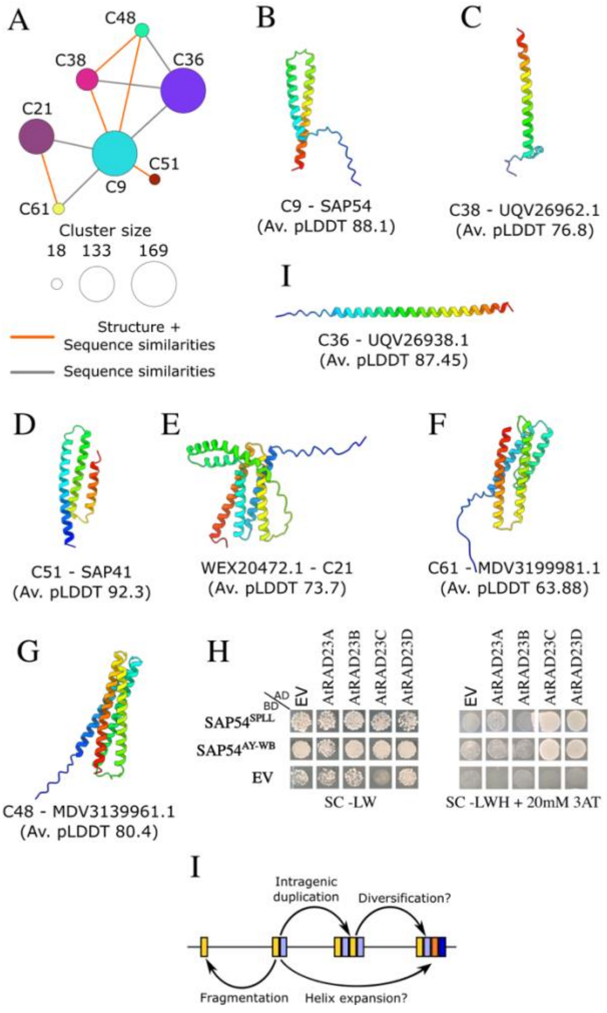


**S13 Figure. Intragenic tandem duplication contributes to the diversification of α-helical folds.** (**A**) PhAME clusters with structural similarities to SAP54 proteins (cluster C9). Node sizes reflect the number of PhAMEs within the clusters. Edges represent pairwise similarities between clusters: grey edges indicate sequence similarities (MMseqs2, E-value ≤ 0.001), and orange edges indicate both structural and sequence similarities (Foldseek and MMseqs2, E-value ≤ 0.001). (**B-G**) Predicted AF2 structures of selected PhAMEs with membership to clusters indicated in (E). Cluster membership and average pLDDT structure confidence scores are shown. (**H**) The C48 cluster SAP54-like protein of Sweet Potato Little Leaf phytoplasma (SPLL) also interacts with *A. thaliana* RAD23 proteins. Y2H assays showing interactions of SAP54$^{AY\text{-}WB}$ and 'SAP54'$^{SPLL}$ with *A. thaliana* RAD23 proteins. Right: Yeast two-hybrid (Y2H) assay to test interactions of SAP54$^{AY\text{-}WB}$ and *A. thaliana* RAD23. EV, empty vector control. AD, GAL4-activation domain. BD, GAL4-DNA binding domain. SD-LWH, triple dropout medium lacking leucine, tryptophan and histidine; 3AT, 3-amino-1,2,4-triazole, a competitive inhibitor of the HIS3 enzyme. Growth on double dropout (–LW) medium confirms the presence of both constructs. (**I**) Schematic overview of evolutionary processes that led to the amplification and diversification of proteins with SAP54-like folds.

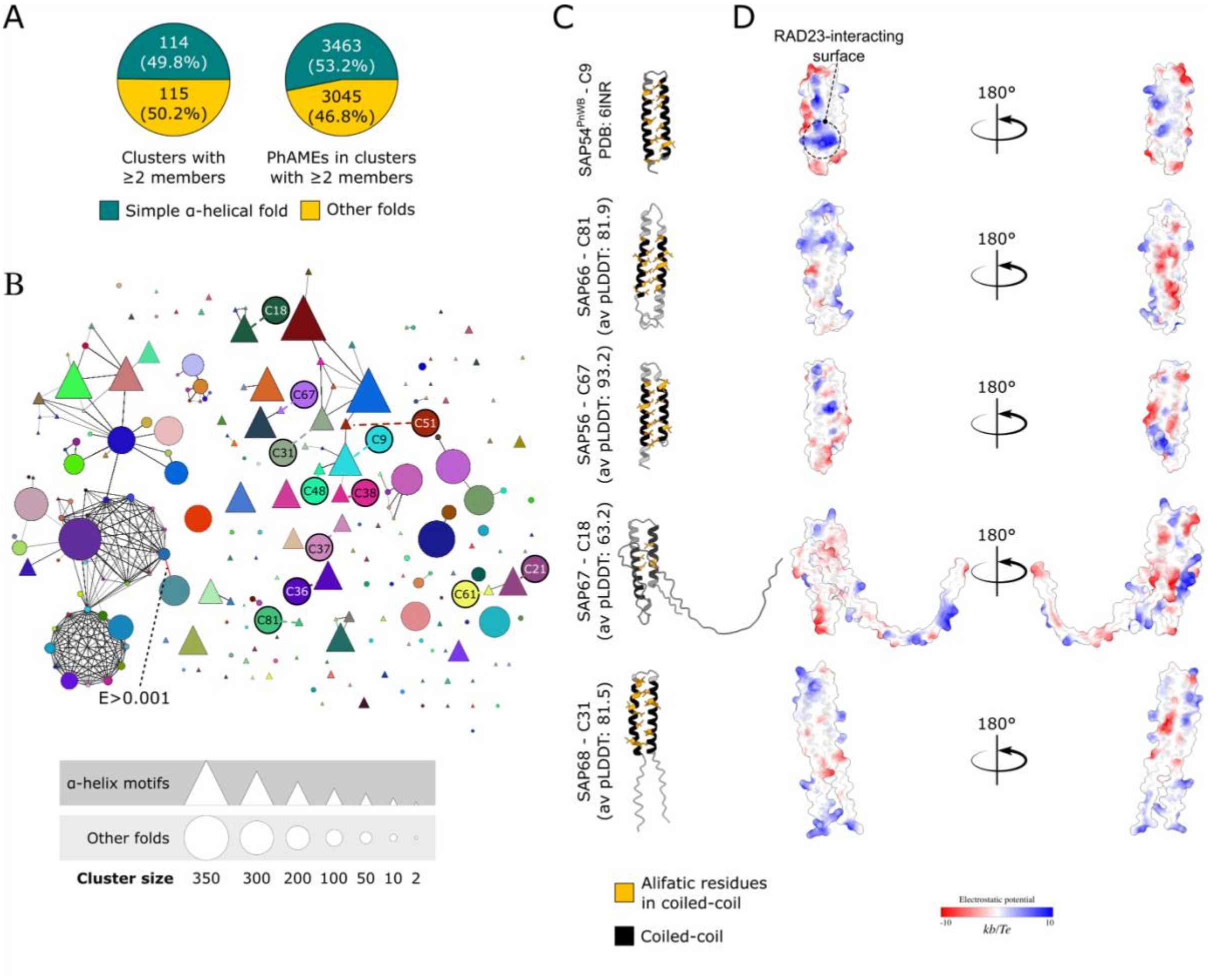


**S14 Figure. PhAMEs consisting of simple α-helical folds contribute to the diversity of the phytoplasma pan-effectorome. (A**) Pie charts showing the proportions of clusters (left) and PhAMEs (right) with small α-helical fold. Singletons were excluded from the analysis. **(B)** Structure similarity network of PhAMEs illustrating relationships among clusters, generated using a more relaxed threshold (Foldseek, E-value ≤ 0.1) than in Fig. 1B, to explore similarities among more divergent folds. Node size is proportional to the number of PhAMEs in each cluster. Clusters predicted to adopt helical structures are shown as triangles, while those with other folds are shown as circles. A single additional structural similarity identified under this relaxed threshold is highlighted in red. Unconnected nodes represent clusters with no detectable structural similarity to others. (**C**) Selected AY-WB phytoplasma effectors composed of antiparallel coiled-coil α-helices, belonging to distinct structural clusters. All effectors share coiled-coil regions predicted with Socket2, shown in black, with hydrophobic faces formed by leucine, valine, and isoleucine residues (shown in orange) oriented toward the protein interior. Cluster membership (as in panel B) and the average pLDDT score for each model are indicated. (**D**) Electrostatic surface potentials of antiparallel coiled-coil α-helix effectors involved in host target binding specificity differ among the effectors shown in panel C. Surfaces are coloured from positive (blue) to negative (red) charge. The positively charged surface of SAP54, involved in interaction with RAD23C, is highlighted. Electrostatic potentials were calculated using the ChimeraX Coulombic command.

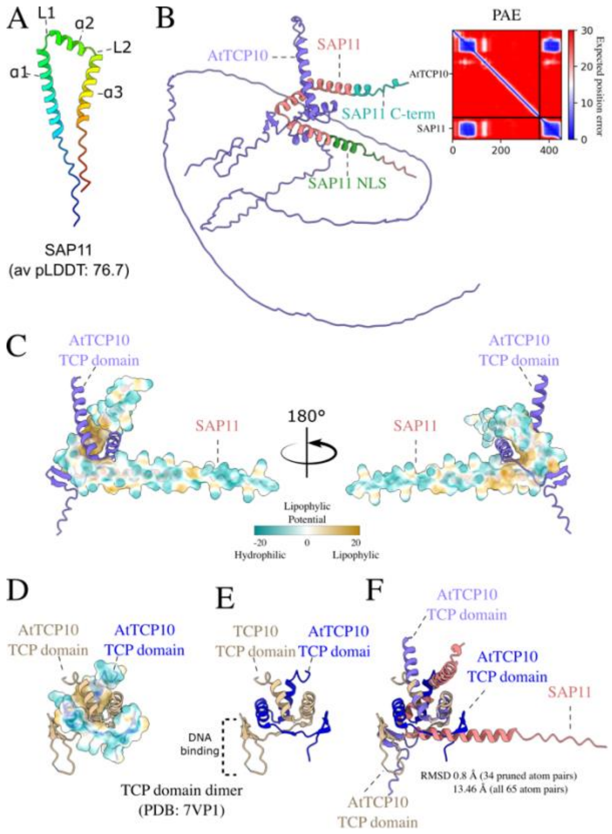


**S15 Figure. Predicted structural features of SAP11 effectors involved in binding plant TCP transcription factors.** (**A**) The AF2-predicted structure of SAP11$^{AY\text{-}WB}$ structure indicating three α-helices (α) and two loops (L). The average pLDDT score is indicated. (**B**) The AF-Multimer-predicted structure of the SAP11$^{AY\text{-}WB}$-AtTCP10 complex. The model predicts that the TCP domain of the transcription factors mediates interaction with SAP11$^{AY\text{-}WB}$, while the SAP11$^{AY\text{-}WB}$ nuclear localization signal (NLS) and C-terminal region are not predicted to contribute to binding, consistent with experimental data (35–37). Predicted Aligned Error (PAE) plot of the highest-confidence model, illustrating the estimated positional uncertainty between residue pairs. Blue indicates low predicted error (high confidence), and red indicates high predicted error (low confidence). (**C**) The pocket created by the two loops, α2 and parts of α1 and α3 creates a lipophilic area that binds the coiled-coil structure of the TCP domain involved in TCP dimerization – this TCP dimerization domain was preciously shown to be required for binding specificity to SAP11 proteins (36). Surfaces are coloured from hydrophilic (dark cyan) to hydrophobic (gold). Lipophilicity potential was calculated using the ChimeraX mlp command. (**D–E**) Experimentally determined structure of a dimer of the coiled-coil region of the TCP10 domain involved in TCP10 dimerization (PDB: 7VPI). In (E), one monomer is shown with lipophilic potential mapped onto its surface representation. DNA binding region is indicated. (**F)** Structural superimposition of the AtTCP10 homodimer and the SAP11–AtTCP10 heterodimer shows that the helix-loop-helix motif of AtTCP10 adopts a similar fold when binding the effector as when interacting with other transcription factors.

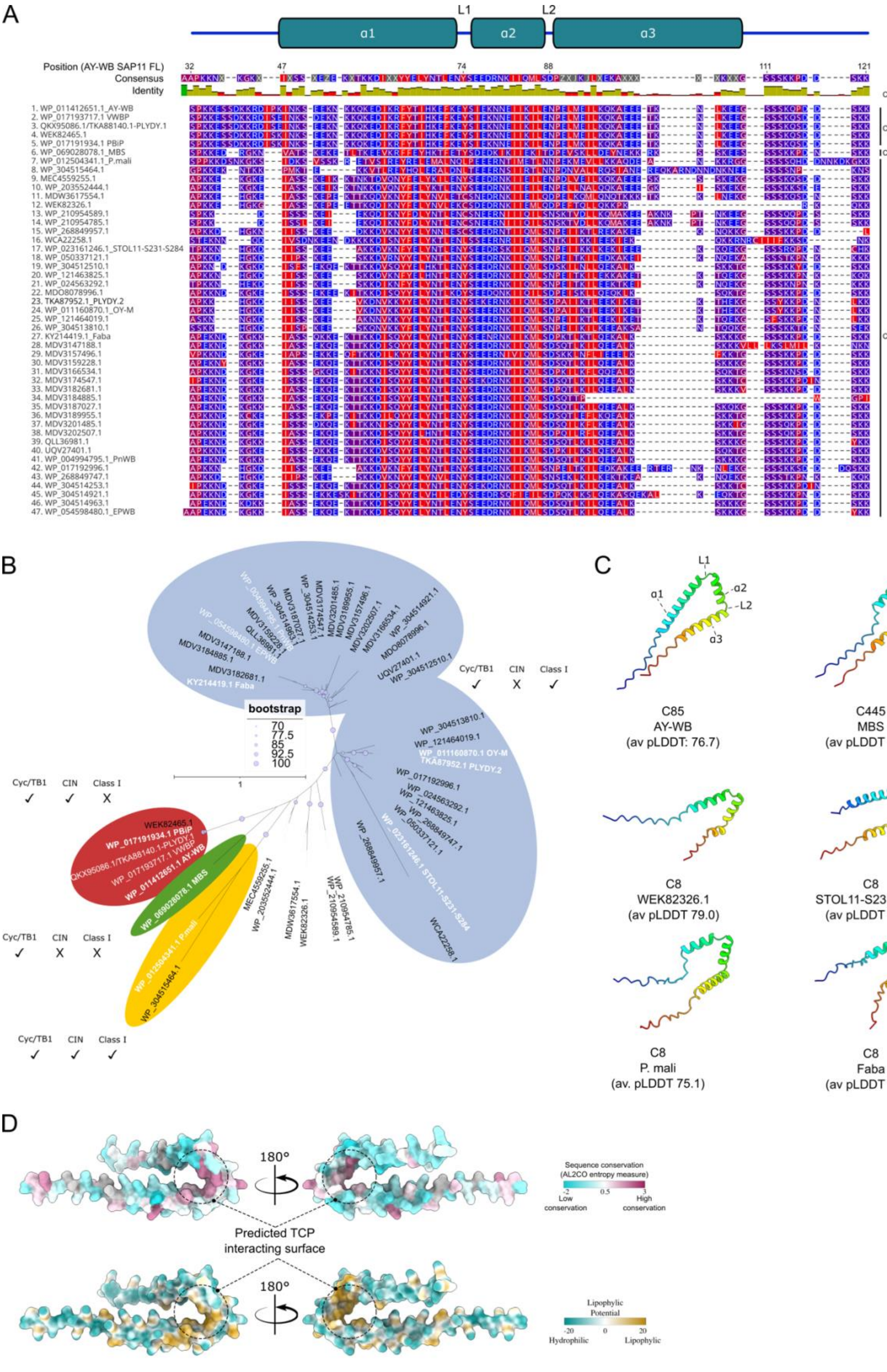
A
L1
L2
α1
α2
α3
Position (AY-WB SAP11 FL)
Consensus
Identity
Cluster
C85
C445
C8
1. WP_011412651.1_AY-WB
2. WP_017193717.1 VWBP
3. QKX95086.1/TKA88140.1-PLYDY.1
4. WEK82465.1
5. WP_017191934.1 PBiP
6. WP_069028078.1_MBS
7. WP_012504341.1_P.mali
8. WP_304515464.1
9. MEC4559255.1
10. WP_203552444.1
11. MDW3617554.1
12. WEK82326.1
13. WP_210954589.1
14. WP_210954785.1
15. WP_268849957.1
16. WCA22258.1
17. WP_023161246.1_STOL11-S231-S284
18. WP_050337121.1
19. WP_304512510.1
20. WP_121463825.1
21. WP_024563292.1
22. MDO8078996.1
23. TKA87952.1_PLYDY.2
24. WP_011160870.1_OY-M
25. WP_121464019.1
26. WP_304513810.1
27. KY214419.1_Faba
28. MDV3147188.1
29. MDV3157496.1
30. MDV3159228.1
31. MDV3166534.1
32. MDV3174547.1
33. MDV3182681.1
34. MDV3184885.1
35. MDV3187027.1
36. MDV3189955.1
37. MDV3201485.1
38. MDV3202507.1
39. QLL36981.1
40. UQV27401.1
41. WP_004994795.1_PnWB
42. WP_017192996.1
43. WP_268849747.1
44. WP_304514253.1
45. WP_304514921.1
46. WP_304514963.1
47. WP_054598480.1_EPWB
B
bootstrap
70
77.5
85
92.5
100
Cyc/TB1 CIN Class I
C
C85
AY-WB
(av pLDDT: 76.7)
C445
MBS
(av pLDDT 76.9)
C8
WEK82326.1
(av pLDDT 79.0)
C8
STOL11-S231-S284
(av pLDDT 78.1)
C8
P. mali
(av. pLDDT 75.1)
C8
Faba
(av pLDDT 77.6)
D
180°
Predicted TCP interacting surface
Sequence conservation (AL2CO entropy measure)
Low conservation
High conservation
Lipophylic Potential
Hydrophilic
Lipophylic

**S16 Figure. The region that mediates binding to the coiled-coil dimerization domain of TCP transcription factors is conserved across SAP11 homologs**. (**A**) MUSCLE (101) protein sequence alignment of the effectors shown in panel A. Secondary structure elements are mapped above the alignment, based on their relative position in the SAP11$^{AY\text{-}WB}$ structure. Cluster membership (C8, C85, C445) is indicated on the right. Residues are coloured by hydrophobicity (136). For effectors with identical sequences a single representative was used for generating the alignment and further analyses. (**B**) RAxML phylogenetic tree constructed using the maximum likelihood method with mature SAP11 and SAP11-like effector sequences from clusters C8, C85, and C445, based on 1,000 bootstrap replicates. Bootstrap values ≥ 70% are indicated by purple circles. White text indicates homologs with confirmed TCP TF binding (73), while black text indicates homologs that have not yet been evaluated. The ability to bind Cyc/TB1, CIN, or Class I TCP transcription factors is indicated as per Correa Marrero et al., 2023 (73). Phylogenetic clades as described in Correa Marrero et al., 2023 are colour-coded. Branch lengths represent amino acid substitution rates (see scale bar). Tree visualisation was performed using the iTOL web server. (**C**) Structural predictions of selected SAP11-related cluster and sub-cluster members, showing a conserved architecture resembling SAP11$^{AY\text{-}WB}$. Cluster membership and average pLDDT scores for each model are indicated. (**D**) Sequence conservation (top) and hydrophilicity (bottom) of SAP11$^{AY\text{-}WB}$ mapped onto its molecular surface. The hydrophobic interface predicted to mediate TCP binding displays high sequence conservation. Sequence conservation and molecular lipophilicity potential were calculated in ChimeraX using AL2CO entropy-based scoring and the mlp command.

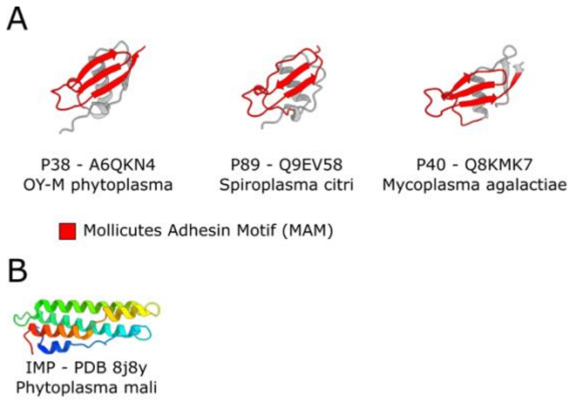


**S17 Figure. The hIG-like domains contain Mollicute adhesin motifs (MAMs).** (A) MAMs mapped within the hIG-like domains (shown in red) of virulence proteins P38, P89, and P40. (B) A MAM was not detected in the experimentally determined structure of the *Ca*. Phytoplasma mali IMP protein (PDB: 8J8Y).
